# dSCOPE: a software to detect sequences critical for liquid-liquid phase separation

**DOI:** 10.1101/2021.01.30.428971

**Authors:** Shihua Li, Kai Yu, Qingfeng Zhang, Zekun Liu, Jia Liu, Huai-Qiang Ju, Zhixiang Zuo, Xiaoxing Li, Zhenlong Wang, Han Cheng, Ze-Xian Liu

## Abstract

Membrane based cells are the fundamental structure and function units of organisms, while evidences were increasing that liquid-liquid phase separation (LLPS) is associated with the formation of membraneless organelles, such as P-bodies, nucleoli and stress granules. Many studies have been undertaken to explore the functions of protein phase separation, but these studies lacked an effective tool to identify the sequence segments that critical for LLPS (SCOPEs). In this study, we presented a novel software called dSCOPE (http://dscope.omicsbio.info) to predict the SCOPEs. To develop the predictor, we curated experimentally identified sequence segments that can drive LLPS from published literature. Then sliding sequence window based physiological, biochemical, structural and coding features were integrated by random forest algorithm to perform prediction. Through rigorous evaluation, dSCOPE was demonstrated to achieve satisfactory performance. Furthermore, large-scale analysis of human proteome based on dSCOPE showed that the predicted SCOPEs enriched various protein post-translational modifications and cancer mutations, and the proteins which contain predicted SCOPEs enriched critical cellular signaling pathways. Taken together, dSCOPE precisely predicted the protein sequence segments critical for LLPS, with various helpful information visualized in the webserver to facilitate LLPS related research.

## INTRODUCTION

Due to the existence of different organelles, eukaryotic cells are divided into different functional domains (1). A growing number of studies have indicated that protein phase separation governs the formation of membraneless organelles, such as nucleoli, stress granules and P bodies, in cells (2–4). In addition, subcellular structures, such as heterochromatin, are also formed by phase separation and have similar potential interactions and physical properties (5,6). When proteins undergo phase separation, they condense into a dense phase that is usually similar to droplets, and the dense phase coexists with the dilute phase (7,8). Phase separation plays critical roles in many biological processes, such as signal transduction, RNA metabolism, and autophagy (8–11). Therefore, to understand the regulation and molecular mechanisms governing phase separation, it is urgently important to identify sequences that are critical for phase separation.

With widespread attention being devoted to phase separation, many databases related to phase separation have been developed. For example, Ning *et al.* (12) collected proteins that are involved in LLPS and constructed an LLPS-associated protein database named DrLLPS, which contains 7,993 scaffolds, 72,300 regulators and 357,594 clients in 164 eukaryotic species. PhaSepDB integrates 2,914 phase separation-associated proteins, and it provides such information as publication source, sequence features and immunofluorescence images of phase separation-associated proteins (13). At present, there are few experimentally verified phase-separated proteins, and the LLPSDB contains only 273 experimentally verified phase-separated proteins (14). The standard of the experimental method to identify the phase-separated structure is that it can form a spherical structure and can be fused. Fluorescence recovery after photobleaching (FRAP) is often performed on droplets as an assessment of their liquidity (15–17). Due to the time-consuming and labor-intensive nature of this method, there is an urgent need to develop an *in silico* method to accurately predict the crucial sequence that drives phase separation, which can facilitate the experimental discovery of phase separation and its mechanism.

In recent years, considerable progress has been made in dissecting the sequence features of proteins that can be phase-separated under physiological conditions. These studies indicate that only certain protein sequences have the ability to undergo phase separation under the proper conditions in living cells. Determining the molecular properties of proteins is crucial to understanding their potential phase behavior. For example, many proteins involved in LLPS have been shown to contain prion-like regions and intrinsically disordered regions. Therefore, the prion-like region prediction tool PLAAC (18) and disorder prediction tools, such as IUPred (19), PONDR-FIT (20) and MobiDB (21), are often used to predict phase separation regions in current research. In addition, hydrophobicity and charged residues are also considered to affect electrostatic interactions, which is further related to phase behavior (22,23). Recently, Vernon *et al.* also compared the prediction performance of these features and pointed out that combining multiple features may facilitate the development of a more accurate prediction tool (24). However, to the best of our knowledge, a tool that can predict the phase separation region by combining multiple features has not been presented to date.

In this study, we built an effective tool for phase separation region detection. First, we manually searched the PubMed literature database and collected all experimentally verified sequences related to phase separation. The final dataset contained 121 phase separation regions in 80 proteins. We intercepted protein sequences through a sliding window of 15 amino acids, and the peptides in the sequences critical for phase separation were defined as positive peptides. Next, eight physiological and biochemical feature scores, including disorder, prion-like, polar, relative surface accessibility (RSA), charge, hydropathy, exposure and low complexity, and four sequence structure features, including amino acid composition (AAC), composition of k-spaced amino acid pairs (CKSAAP), position-specific scoring matrix (PSSM) and binary encoding profiles (BE), were extracted and modeled with a random forest algorithm. The TOPT package was utilized to adjust the hyperparameters and to optimize the prediction model. By 4-, 6-, 8-, and 10-fold cross-validation in the training dataset, dSCOPE showed excellent robustness and satisfactory performance. In addition, by utilizing dSCOPE, we comprehensively analyzed the relationship between SCOPEs and tumor mutations, as well as posttranslational modifications, which may provide helpful information for the diagnosis and treatment of some diseases.

## MATERIALS AND METHODS

### Data collection and preparation

To collect the experimentally identified sequences that can drive protein phase separation, we employed the keyword “phase separation” to retrieve the published literature from PubMed (http://www.ncbi.nlm.nih.gov/pubmed), and eventually, we collected 121 sequences from 80 proteins that were essential for phase separation. The sequence of each protein was retrieved from the UniProt database (25). We treated all regions that play a crucial role in phase separation as positive sequences, and other fragments of the same protein were treated as the negative sequences. For each dataset, we generated 15-length peptides through the sliding window with a step size of 8. Among these peptides, the peptides in human proteins were treated as the training dataset, while the peptides in yeast proteins were treated as the testing dataset. In total, we obtained 1,737 positive peptides and 3,125 negative peptides for the training dataset and 379 positive peptides and 1,075 negative peptides for the testing dataset.

### Feature extraction

#### Physicochemical properties

In recent years, researchers have made considerable progress in analyzing the sequence characteristics of proteins that can undergo phase separation under physiological conditions. The common feature of LLPS proteins is the presence of an intrinsic disordered region (IDR) with multiple interacting motifs (16,26). Charge pattern, amino acid composition, and solubility also affect phase separation (27). We calculated the disorder scores of the proteins by IUPred (19), the per-residue prion-like scores were obtained from PLAAC (18), exposure and surface accessibility analysis were performed by NetSurfP (28), the hydrophobicity was based on the theory of Kyte, J *et al*. (29), as well as charge from Fauchere *et al.* (30), and we used StatSEG (https://github.com/jszym/StatSEG) to obtain the low-complexity region scores. In addition, we also considered the polarity of amino acids.

#### Composition of k-spaced amino acid pairs (CKSAAP)

Similar to Zhao *et al*. (31), we used the composition ratio of residue pairs of k intervals in the protein sequence fragments in the sequence to establish a mathematical model and extract feature vectors. In other words, if a peptide consists of 20 kinds of amino acids, each amino acid and its next adjacent amino acid form a pair of extracted amino acids, that is, the separation distance between these two amino acids is k = 0 amino acids, then there are 400 possible amino acid pairs (e.g., AA, AC, AD, and so on). According to the probability of these residue pairs appearing in this protein sequence, a 441-dimensional feature vector is generated. With the increase in the k value, although the accuracy and sensitivity of the prediction model increases, the calculation time and cost of the random forest model training also increases notably. In this regard, only the CKSAAP coding with k values equal to 0, 1, 2, and 3 are considered; therefore, the total dimension of the feature vector is 400 × 4 = 1,600.

#### Position-specific scoring matrix (PSSM)

PSSM is a common feature extraction method in biological sequence analysis, also known as the position weight matrix (32,33). This matrix has 20 × M elements, where M is the length of the target sequence. The occurrence frequency of different amino acids at each position in the matrix was calculated, and the details are as follows:

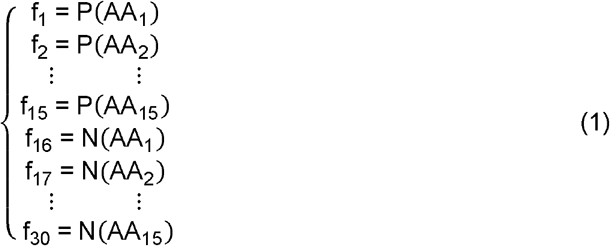

In Eq. 1, the peptides consist of 15 amino acids; P(X1) represents the occurrence frequency of amino acid AA1 at position 1 in the positive group, while N(X1) denotes the occurrence frequency of amino acid AA1 at position 1 in the negative group. Therefore, each peptide can be represented by a position weight amino acid composition vector with dimensions of 30.

#### Amino acid composition (AAC)

AAC is an elementary feature and describes the frequency of occurrence of each amino acid in the sequence (34). The dimension of AAC is 20 in this work.

##### Binary encoding profiles (BE)

Binary encoding is similar to the binary language of computers (35). We converted each sequence into a combination of 20-dimensional vectors. For example, if a sequence is ARDCQEHIGNLKMFPSTWYV, then amino acid A corresponds to (10000000000000000000), and amino acid V corresponds to (00000000000000000001). In this work, the vector size is 300.

### Machine learning classifiers

To predict the sequence that is important to phase separation, the random forest algorithm is introduced into our prediction. For a given protein sequence, a short peptide consisting of 15 amino acids in length is intercepted through a sliding window. Each adjacent short peptide needs to be separated by 8 amino acid residues, and short peptides less than 15 in length are removed. Protein fragments are encoded by eight physical and chemical features scores and four feature extraction methods. Next, we tested the performance of five ML classifiers, including logistic regression, random forest, LDA, AdaBoost, and KNN, then adopted the final algorithm for prediction based on the performance. Moreover, to generate the optimal performance, the TOPT (https://github.com/EpistasisLab/tpot) package was integrated to optimize the hyperparameters.

### Performance evaluation

We used the following key measurements to evaluate the performance of the model: specificity (*Sp*), sensitivity (*Sn*), accuracy (*Ac*), and receiver operating characteristic (ROC) curves were drawn, and AUC (area under ROC) values were calculated. The measurements were defined as follows:

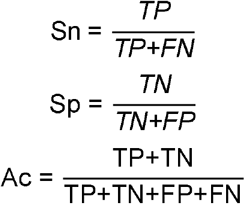

In this work, 5- and 10-fold cross-validation was performed. In addition, an independent testing set was used to prove the advantages of our model compared to the current commonly used tools.

### Implement of the webserver

We constructed the dSCOPE web server in Python, PHP, JavaScript and HTML, which can be freely accessed at http://dscope.omicsbio.info. NetSurfP was adopted to predict the surface accessibility and secondary structure information for each sequence, the query protein disorder information was calculated by IUPred (19), the per-residue prion-like scores were obtained from PLAAC (21), and the charge pattern from Fauchere *et al.* (30), amino acid composition, and solubility also affect phase separation (27). Additionally, the hydrophobicity was determined based on the theory of Kyte, J *et al*. (29), and subcellular location was obtained from UniProt.

### Molecular condensate enrichment analysis

We used dSCOPE to predict 3,633 human proteins related to phase separation retrieved from DrLLPS and selected 998 proteins that contain the potential SCOPEs. Then, we performed molecular condensate enrichment analysis with the 998 proteins, and Fisher's exact test was used to test the significance. We further used WocEA to display the result in the word cloud (36).

### Kinase and TF enrichment analysis

Kinase- and phosphorylation-related information was collected and curated from multiple phosphorylation databases verified by a number of experiments, including dbPTM (37), BIOGRID (38), PHOSIDA (39), phosphoELM (40), PhosphositePlus (41), and RegPhos2.0 (42), which included 192,111 phosphorylation sites. Next, we used Fisher's exact test to perform kinase enrichment analysis on potential phase-separated proteins. In addition, transcription factor (TF) enrichment analysis used ChEA3 (43).

### Pathway enrichment analysis

To better understand the potential function of phase separation proteins, we used the R ClusterProfiler package for Gene Ontology (GO) function annotation (44), and the enrichment analysis of these proteins in Kyoto Encyclopedia of Genes and Genomes (KEGG) pathways was performed using KOBAS (45). The visible network was constructed using Cytoscape (46).

### Mutation analysis of phase separation-related regions

We downloaded GDC TCGA somatic mutations from 11 cancer types (BLCA, BRCA, CESC, COAD, HNSC, LIHC, LUAD, LUSC, SKCM, STAD, UCEC) from the Xena Browser (https://xenabrowser.net/datapages/), removed redundant mutations, and retained only missense mutations for further analysis. We generated peptide windows composed of 15 amino acids. Each region in a protein would generate two peptides: one was extracted from the origin sequence, and the other was from the new sequence obtained after mutation. Finally, we predicted the probability for these sequence windows before and after mutation based on dSCOPE. Based on the prediction result of dSCOPE, we further divided the mutation into two types according to whether the SCOPEs were affected.

## RESULTS

### Construction of the computational model to predict the crucial region of LLPS

We used the keyword “phase separation” to manually collect experimentally confirmed phase-separated proteins from the literature (Figure 1), and we obtained the training and testing datasets as mentioned above. We eventually obtained 1,737 positive sequences and 3,125 negative sequences for *Homo sapiens* as the training dataset and 379 positive sequences and 1,075 negative sequences for *Saccharomyces cerevisiae* as the testing dataset. Then, we developed dSCOPE software for detecting sequences critical for phase separation-related proteins based on random forest and TPOT algorithm. Eight physicochemical property (PP) scores of the sequences were extracted, including disorder, exposure, polar, low complexity region, charge, prion-like region, surface accessibility, and hydropathy. In addition, the sequence features, including AAC, CKSAAP, PSSM, and BE, were also considered (Figure 1). Furthermore, we used Python, PHP, JavaScript and HTML to construct the dSCOPE online server, which can be accessed through http://dscope.omicsbio.info. Finally, a series of further analyses were performed using dSCOPE, including molecular condensates, posttranslational modification (PTM) analysis of potential SCOPEs, GO functional annotation, KEGG pathway, kinase, and transcription factor enrichment (Figure 1).

**Figure 1.**
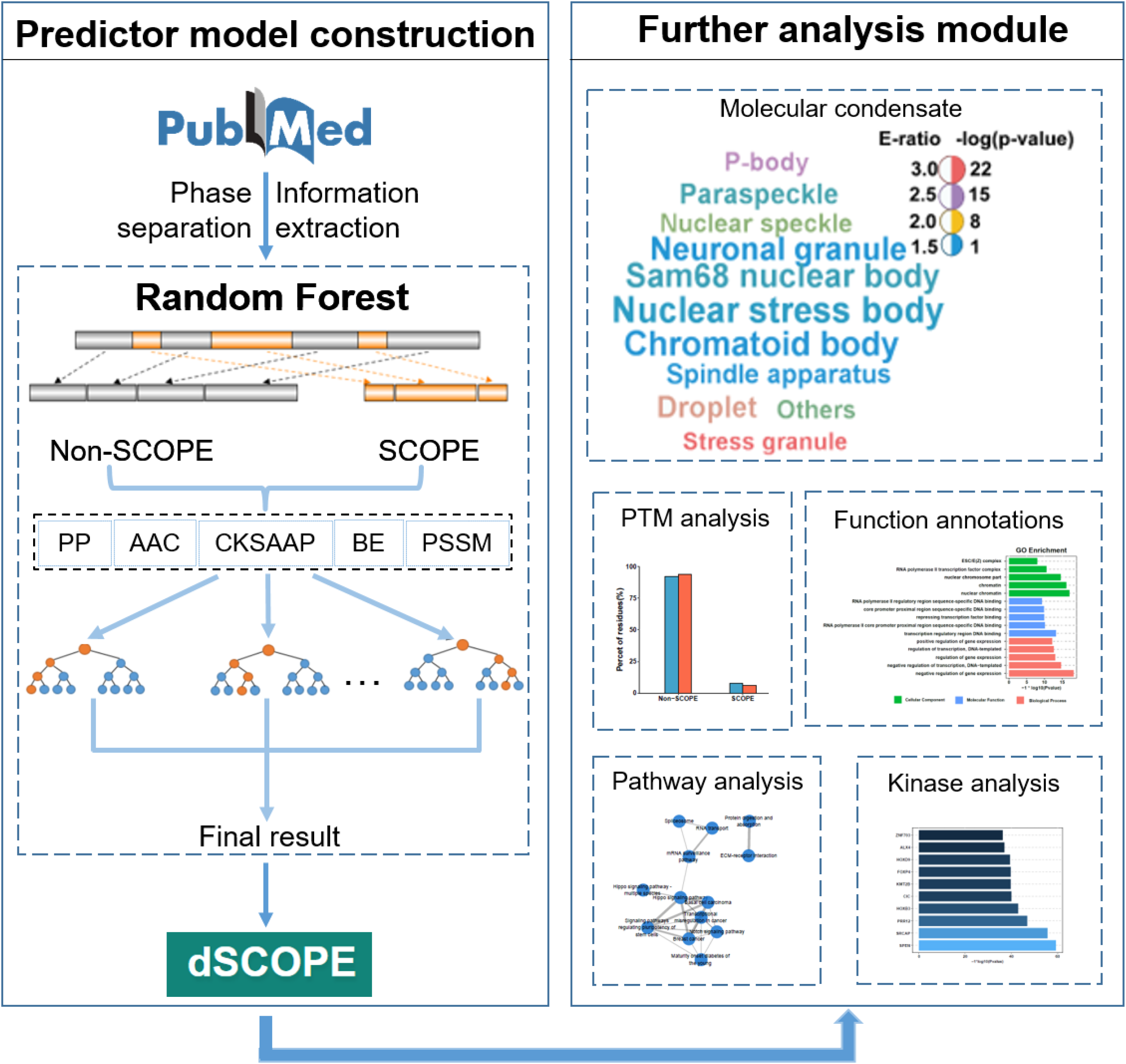
Overview of the model. The key regulatory sequence of phase separation was extracted from PubMed, and then dSCOPE was constructed by using random forest development. Downstream analysis reveals the characteristics of phase separation.

### Evaluating the performance of dSCOPE

We compared the eight reported features related to LLPS between SCOPEs and non-SCOPEs (Figure 2A). This comparison indicated that these features are significantly different in SCOPEs and non-SCOPEs. The SCOPE shows more disorder, the sequence net charge approaches zero, the complexity is low, and the hydrophilic polar amino acids are enriched. This result is consistent with the findings of a previous report (47). In the analysis of amino acid frequency, glycine (G), asparagine (N), proline (P), glutamine (Q), serine (S) and tyrosine (Y) appeared more frequently in the key regulatory sequence of phase separation (Figure 2B), and most of these amino acids are polar amino acids.

**Figure 2.**
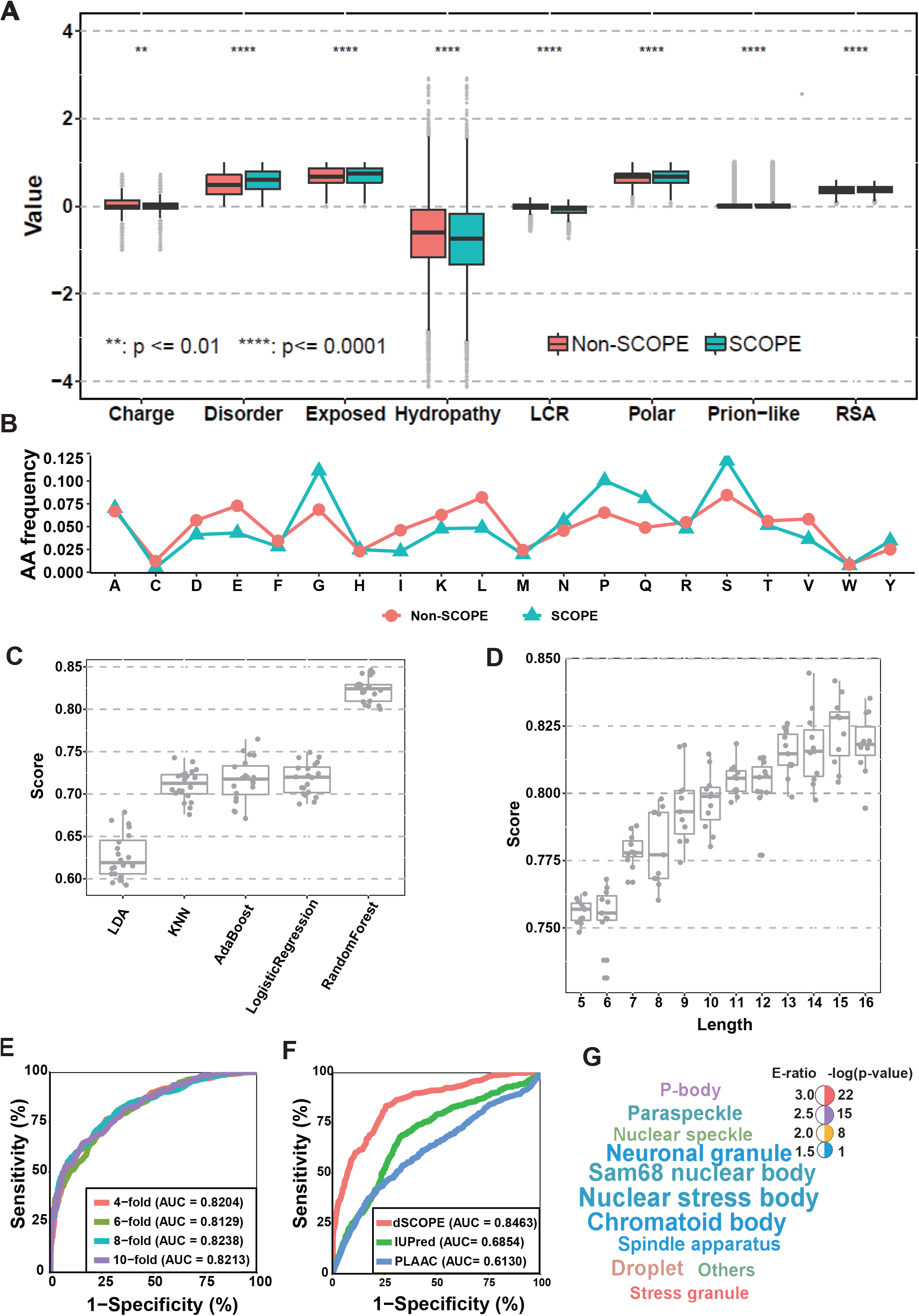
Features of the phase separation region and performance of dSCOPE. (A) Features of the phase separation regions. (B). The frequency of amino acids in different regions. (C) AUC values of different lengths. (D) Use 5-fold cross validation to compare the performance of machine learning algorithms; the median values are 0.8242 (Random forest), 0.7199 (Logistic regression), 0.7177 (AdaBoost), 0.7127 (KNN) and 0.6191 (LDA). (E) The 4-, 6-, 8-, and 10-fold cross validation results in the training dataset. (F) Comparison of the models with other tools. (G) DrLLPS enrichment analysis.

Based on the model architecture described in the method, we compared five machine learning algorithms through 5-fold cross-validation. As shown in Figure 2C, the random forest algorithm outperformed the other ML algorithms, and it has higher robustness and faster calculation speed. In addition, we intercepted different lengths and used 5-fold cross-validation to evaluate its performance. When the length increased, the AUC value gradually increased, but when the length reached 16, it decreased instead (Figure 2D). Therefore, we chose a length of 15 to ensure prediction performance. Then, we developed dSCOPE software for detecting sequences critical for phase separation-related proteins based on random forest algorithm, and the hyperparameters were optimized by the TPOT package. 4-, 6-, 8- and 10-fold cross-validation were employed to evaluate the prediction performance of dSCOPE. The ROC curves were shown and AUCs were calculated (Figure 2E). As shown in Figure 2E, the AUC values were 0.8204 (4-fold), 0.8129 (6-fold), 0.8238 (8-fold), and 0.8213 (10-fold). The results of the various validations were very similar to one another, which indicated that dSCOPE is a stable and robust predictor.

To verify the superiority of the dSCOPE model, we compared it with two widely used phase-separated region detection tools, IUPred (19) and PLAAC (18), even though they were not developed for phase separation prediction. We evaluated the prediction performances of these tools using the testing dataset. The AUC values for dSCOPE were 0.8463, while those for IUPred and PLAAC were 0.6854 and 0.6130 (Figure 2F), respectively. In summary, our prediction tool achieved good prediction performance.

### Development and application of dSCOPE

We developed a freely available website for phase separation prediction to facilitate scientific research. The dSCOPE was implemented in Python, PHP, JavaScript and HTML, and the prediction page is shown in Figure 3A. Users should paste the FASTA format sequences of their proteins of interest into the text box and then choose an organism and a threshold to obtain the result. We also provided some examples to show the usage of dSCOPE; meanwhile, users only need to click the “Example” button to see the default protein sequences and their predicted results. Users can also use UniProt ID, gene name or protein name in the search interface to directly determine the predicted results of human protein (Figure 3B). On the results page, the prediction information was organized into three sections, including “dSCOPE prediction results”, “Information of the protein” and “Sequence and structural characteristics of the protein” (Figure 3C-D). The “dSCOPE prediction results” was a detailed description of the prediction results (Figure 3D). If the users wanted to search for predicted results regarding a reviewed human protein by UniProt ID, gene name or protein name, the protein details annotated by UniProt were displayed in “Information of the protein” (Figure 3C). The “Sequence and structural characteristics of the protein” included the protein predicted potential phase separation regions, the dSCOPE score for each residue, the sequence and structure properties, and the protein subcellular location information (Figure 3D). Supporting multiple protein sequence predictions, users can select the protein prediction information to display by clicking on the selection box at the top. In conclusion, dSCOPE appears to be a comprehensive web server for protein phase separation prediction to facilitate related research.

**Figure 3.**
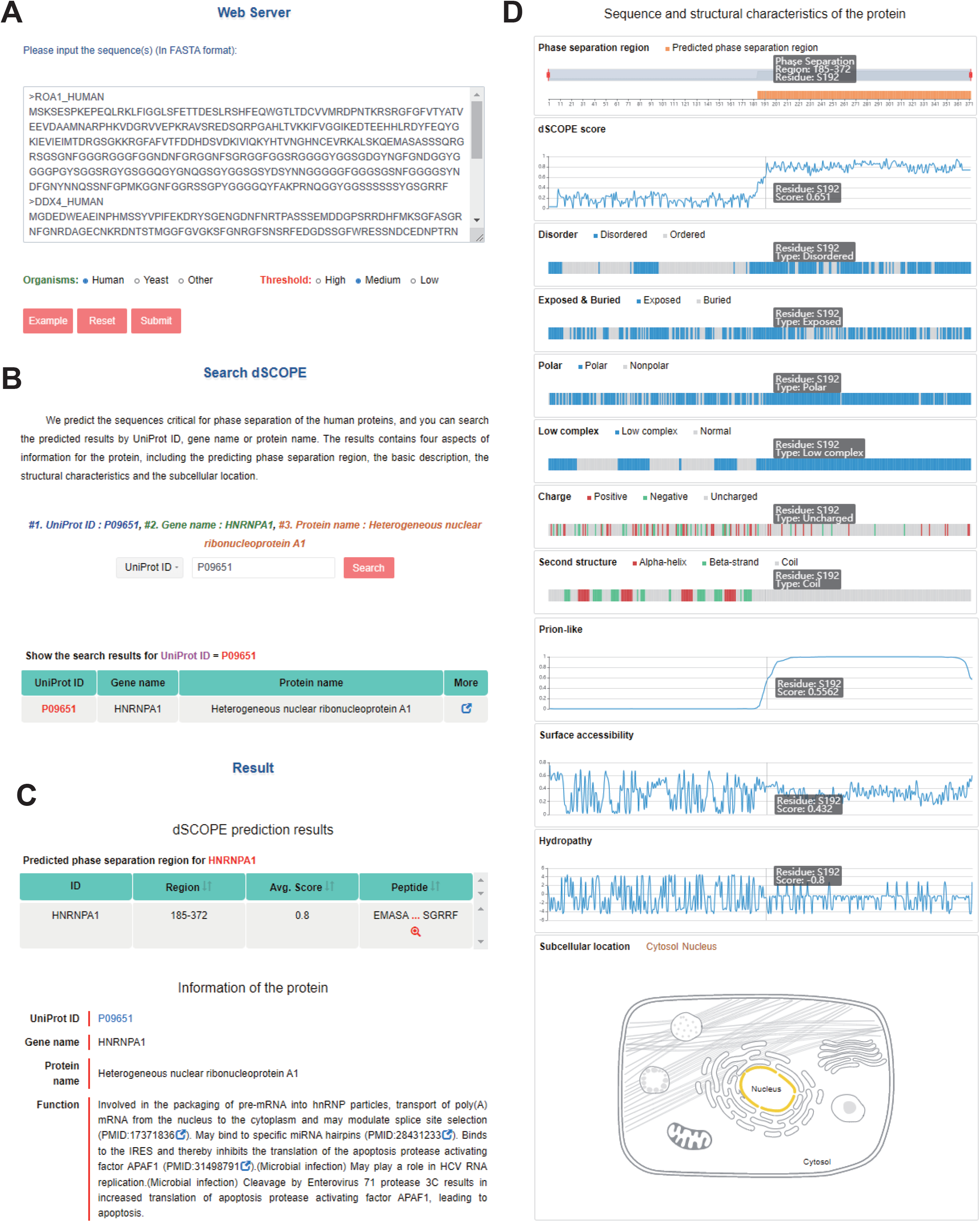
Web server of dSCOPE. (A) The prediction page. (B) The search page. (C) Potential phase separation sequences. (D) The sequence and structure properties of the query protein and subcellular location.

To better characterize the application of dSCOPE, we predicted 3,633 human proteins related to phase separation retrieved from DrLLPS and selected 998 proteins containing the potential SCOPE. The prediction results showed that proteins potentially containing sequences that are critical for phase separation are significantly enriched in the P-bodies, stress granules, and nuclear speckles (Figure 2G). Previous studies have also shown that phase-separated proteins primarily exist in membraneless organelles, such as P-bodies and stress granules.

### Functional analysis of potential phase separation proteins

In phase-separated proteins, the sequences critical for phase separation possess numerous disordered regions, and these regions are susceptible to various PTMs (48). To further investigate whether the protein containing SCOPE extracted from dSCOPE has the potential for phase separation, we predicted 20,380 reviewed human proteins, and under the condition of FDR<1%, 4,637 proteins containing potential sequences essential for phase separation were selected. Therefore, we employed the data of PLMD (http://plmd.biocuckoo.org/) and EPSD (http://epsd.biocuckoo.cn/) to analyze and understand the relationship between SCOPE and PTMs. Lysine modification usually was evenly distributed between protein sequences but was slightly enriched in the potential SCOPE (E-ratio = 1.40, *p-value* = 5.04 × 10^−12^, Fisher’s exact test) (Figure 4A), and the phosphorylation sites were also enriched in the SCOPE (E-ratio = 1.25, *p-value* = 1.17 × 10^−123^, Fisher's exact test) (Figure 4B).

**Figure 4.**
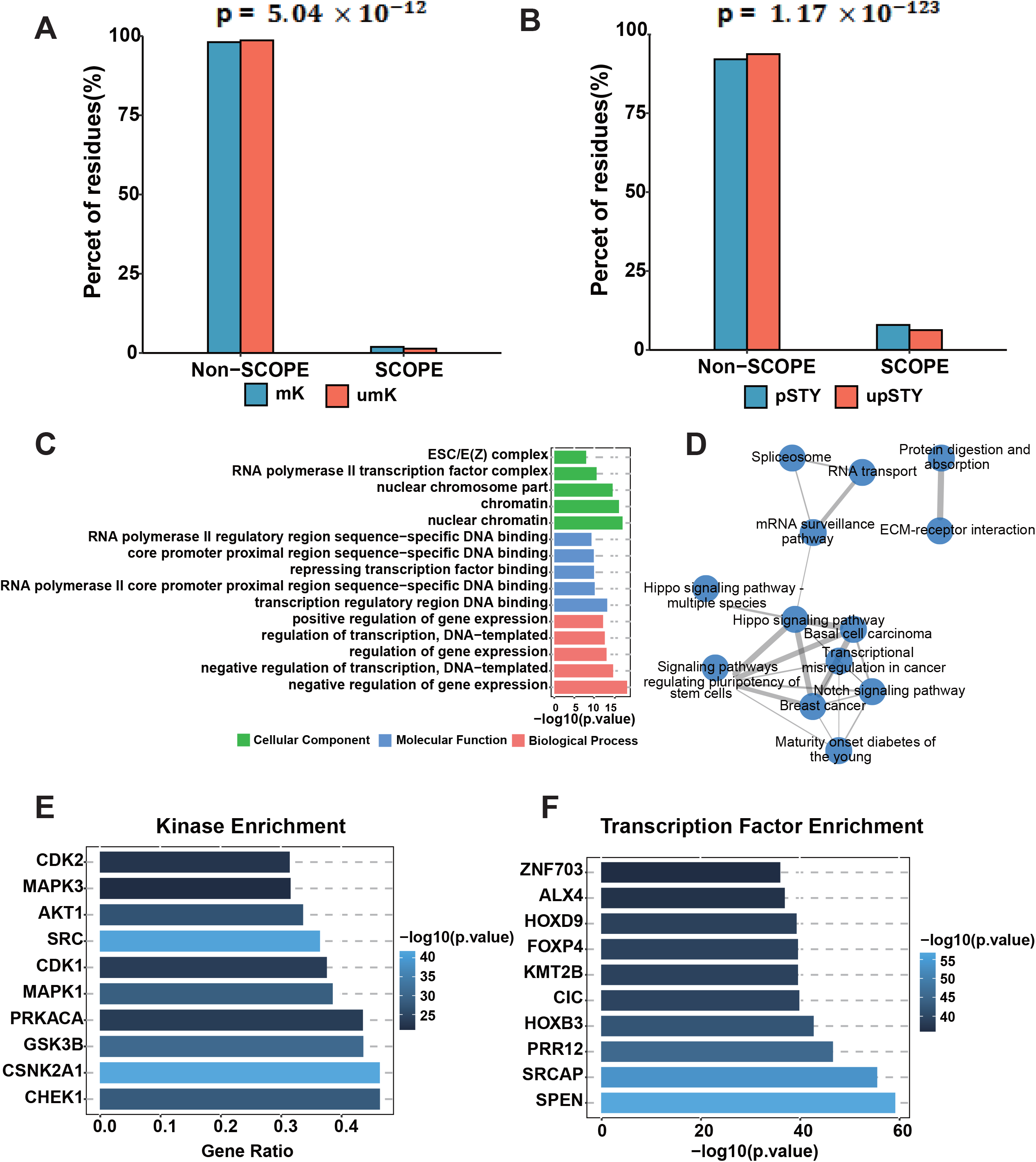
Functional analysis of SCOPEs and potential phase separation proteins. (A) Lysine modification enrichment. mK, modified lysine residue; umK, unmodified lysine residue. (B) Phosphorylation enrichment. pSTY, phosphorylated STY residue; upSTY, unphosphorylated STY residue. (C) GO enrichment. (D) KEGG pathway interaction network. (E) Enrichment of upstream kinases for phosphoproteins in potential phase separation proteins. (F) TF enrichment analysis.

Not only that, we also performed GO functional annotation. As shown in Figure 4C, the proteins are highly enriched in chromatin and various protein complexes. In addition, many proteins may be involved in transcription and may be related to gene expression and regulation. KEGG pathway enrichment analysis revealed that potential phase-separated proteins are significantly enriched in several pathways related to proliferation and apoptosis, such as the Hippo signaling pathway, the Notch signaling pathway and signaling pathways regulating pluripotency of stem cells (Figure 4D). It can be observed from previous studies that these pathways contain various posttranslational modification processes, especially phosphorylation (49–51).

Among the 4,637 proteins with potential SCOPE, 3,099 proteins have phosphorylation sites. In addition, 841 proteins have phosphorylation sites for known kinases. We used Fisher’s exact test to perform kinase enrichment analysis on phosphoproteins containing known kinases. As shown in Figure 4E, we found that MAPK1, SRC, CDK1, and CDK2 were significantly enriched. SRC has been shown to play a key role in the phase separation of the FUS and tau proteins (52,53). In addition, phase-separated proteins have been extensively confirmed to be involved in gene expression and regulation, and identifying transcription factors that lead to changes in gene expression is an important step in elucidating gene regulation networks (54). Therefore, we performed a transcription factor enrichment analysis using ChEA3 (43). The enrichment results are shown in Figure 4F. The top 5 TFs were SPEN, SRCAP, PRR12, HOXB3, and CIC. In these enriched TFs, CIC usually cooperates with ATXN1 to play a role in the development of the central nervous system, and ATXN1 is a proven phase separation protein(55). In addition, SPEN also regulates the notch signaling pathway (56).

## DISCUSSION

In recent years, research on the mechanism of phase separation of biomolecules has increased rapidly. Accumulating evidence demonstrates that LLPS underlies the formation of membraneless organelles (2). Furthermore, LLPS is also related to various neurodegenerative diseases, such as amyotrophic lateral sclerosis (ALS) (57,58), Alzheimer’s disease (AD) (59) and frontotemporal dementia (FTD) (60). However, the prediction of protein LLPS through computational methods remains a challenge. PLAAC developed with hidden Markov is a tool that is often used to predict phase separation, but its initial development was to identify prion-like sequences (18). Vernon *et al.* compared the prediction performance of six sequence-based algorithms and one empirical approach that has recently been employed to predict protein phase separation (24). It was observed that considering multiple features may help researchers obtain more accurate predictions. Hence, we developed dSCOPE in consideration of various features.

Most sequences critical for phase separation contain IDRs, which cause them lack secondary structures and therefore be particularly susceptible to PTMs, especially phosphorylation and lysine modification (48,61). This property was also demonstrated by the analysis of lysine and phosphorylation modifications. And in order to explore the relationship between cancer mutations and phase separation, we conducted a mutation analysis on the collected phase-separated proteins and proteins predicted by dSCOPE and found that mutations are more likely to occur in non-SCOPE phase-separated proteins (Supplementary Figure 1A-B).

Previous studies have shown that phase separation plays an important role in chromosome structure (62). This role is consistent with our analysis of potential phase separation proteins. In addition, KEGG enrichment analysis of potential phase-separated proteins showed that pathways related to proliferation, apoptosis and signal transduction processes had significant enrichment. Increasing experimental evidence shows that phase separation is involved in the process of cell autophagy (63). The functional preference of these predicted proteins is consistent with the characteristics of experimentally confirmed phase-separated proteins, indicating that our tool is reliable, and these analyses may provide experimental researchers with some new ideas.

dSCOPE is the first tool to combine multiple features to predict the sequences that are critical for phase separation. Various feature scores, including disorder, prion-like, polar, RSA, charge, hydropathy, exposure and low complexity, are examined. Although dSCOPE has achieved good prediction performance, there are many aspects that warrant improvement. It is well-known that a larger training dataset produces more accurate predictive performance. In the future, experimentally identified sequences with phase separation proteins will be continuously collected from the literature and integrated into the predictive model when available. Furthermore, more machine learning algorithms need to be considered, such as deep neural networks (DNNs) and recurrent neural networks (RNNs), which may also improve the current prediction performance.

## DATA AVAILABILITY

The dSCOPE server is freely available at http://dscope.omicsbio.info.

## ABBREVIATIONS

LLPS: liquid-liquid phase separation
SCOPE: sequence critical for phase separation
ML: machine learning
RNN: recurrent neural network
DNN: deep neural network
LDA: Linear Discriminate Analysis
KNN: K-Nearest Neighbor
PP: physicochemical properties
CKSAAP: composition of k-spaced amino acid pairs
BE: binary encoding profiles
PSSM: position-specific scoring matrix
AAC: amino acid composition
FDR: false discovery rate
Sp: specificity
Sn: sensitivity
Ac: accuracy
ROC: receiver operating characteristic
AUC: area under ROC curve
RSA: relative surface accessibility
TF: transcription factor
mK: modified lysine residue
umK: unmodified lysine residue
pSTY: phosphorylated STY residue
upSTY: unphosphorylated STY residue

## FUNDING

This work was supported by grants from the Natural Science Foundation of China (91953123 to ZX.L., 31601067 to H.C.); Program for Guangdong Introducing Innovative and Entrepreneurial Teams (2017ZT07S096 to ZX.L.); Pearl River S&T Nova Program of Guangzhou (201906010088 to ZX.L.); Tip-Top Scientific and Technical Innovative Youth Talents of Guangdong Special Support Program (2019TQ05Y351 to ZX.L.); and Key program for Department of Science and Technology of Qinghai province (2017-ZJ-Y13 to H.C.).

## CONFLICT OF INTEREST

The authors declare no competing financial interests.

## FIGURE LEGENDS

**Supplementary Figure S1.** Enrichment analysis of tumor mutations. (A) Tumor mutations in SCOPEs confirmed by experiment. (B) Tumor mutations in potential SCOPEs.

## Notes

http://dscope.omicsbio.info

### Competing Interest Statement

The authors have declared no competing interest.

