## Supplementary Data for "dSCOPE: a software to detect sequences critical for liquid-liquid phase separation"

**Supplementary Figure S1. Enrichment analysis of tumor mutations.** (A) Tumor mutations in SCOPE confirmed by experiment. (B) Tumor mutations in potential SCOPE.

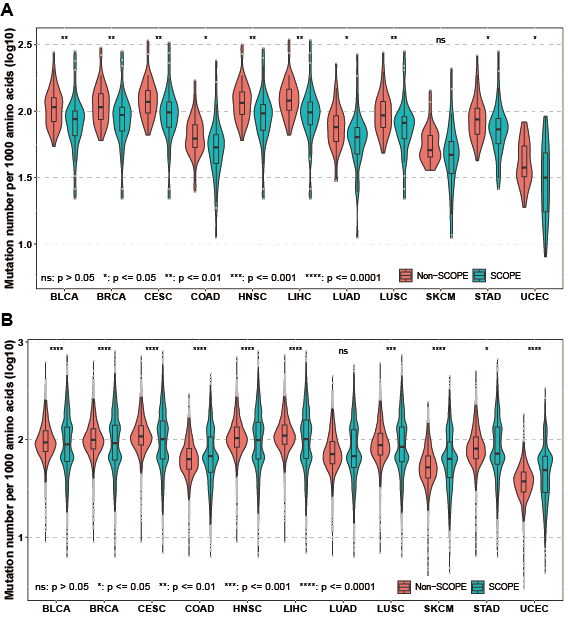
