## Supplementary figures and images for "dSCOPE: a software to detect sequences critical for liquid-liquid phase separation"

### Supplementary Figure S1

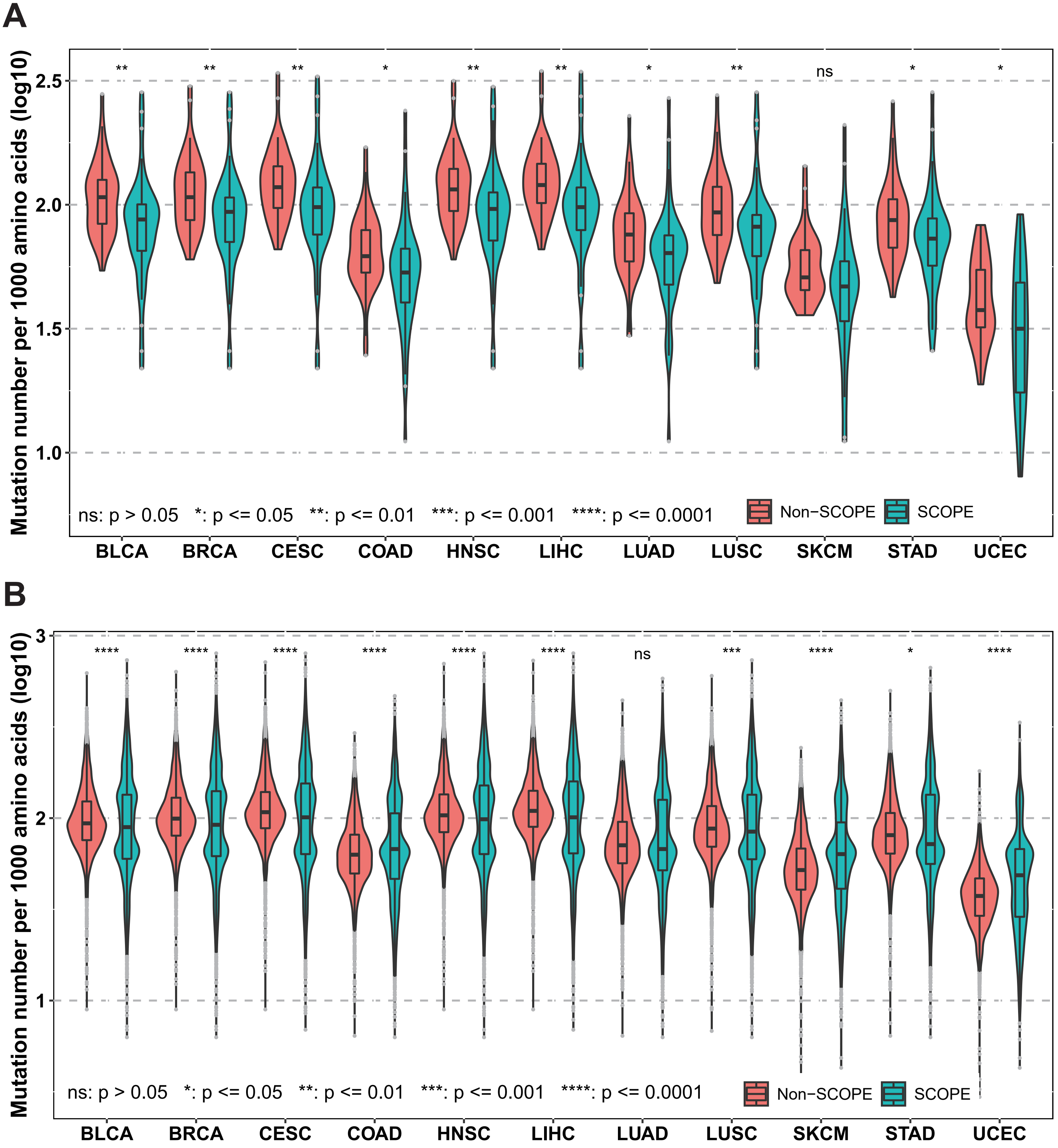
